# CNP blocks mitochondrial depolarization and inhibits SARS-CoV-2 replication *in vitro* and *in vivo*

**DOI:** 10.1101/2023.06.09.544327

**Authors:** James Logue, Victoria M. Melville, Jeremy Ardanuy, Matthew B. Frieman

## Abstract

The COVID-19 pandemic has claimed over 6.5 million lives worldwide and continues to have lasting impacts on the world’s healthcare and economic systems. Several approved and emergency authorized therapeutics that inhibit early stages of the virus replication cycle have been developed however, effective late-stage therapeutical targets have yet to be identified. To that end, our lab identified that 2’,3’ cyclic-nucleotide 3’-phosphodiesterase (CNP) inhibits SARS-CoV-2 virion assembly. We show that CNP inhibits the generation of new SARS-CoV-2 virions, reducing intracellular titers without inhibiting viral structural protein translation. Additionally, we show that targeting of CNP to mitochondria is necessary for inhibition, blocking mitochondrial depolarization and implicating CNP’s proposed role as an inhibitor of the mitochondrial permeabilization transition pore (mPTP) as the mechanism of virion assembly inhibition. We also demonstrate that an adenovirus expressing virus expressing both human ACE2 and CNP inhibits SARS-CoV-2 titers to undetectable levels in lungs of mice. Collectively, this work shows the potential of CNP to be a new SARS-CoV-2 antiviral target.

**Author Summary:** Upon entry into a cell, viruses manipulate the cellular environment for the benefit of their replication. We have found that upon infection, SARS-CoV-2 induces the mitochondria to release reactive oxygen species (ROS) into the cytoplasm which benefits the replication of the virus. We identified a phosphodiesterase, called CNP, that blocks this release via inhibition of an inducible pore called the mitochondrial permeability transition pore or mPTP. We also found that a drug targeting this pore inhibits SARS-CoV-2 replication. We test the function of CNP in vivo and find that if we overexpress CNP in mouse lungs, we inhibit SARS-CoV-2 replication. Together this demonstrates a key function of the mitochondria on SARS-CoV-2 replication and that antithetically ROS release enhances viral replication. We propose this is common across coronaviruses and potentially other viruses identifying a novel target for future therapies.

## Introduction

Severe acute respiratory syndrome coronavirus 2 (SARS-CoV-2) is the etiologic agent of coronavirus disease 2019 (COVID-19), first reported in Wuhan City, Hubei Provence, China in December 2019 [1–3]. Since then, SARS-CoV-2 infections have expanded to a worldwide pandemic, with over 675 million reported cases and over 6.8 million reported deaths at the time of writing [4, 5]. In the absence of broad-spectrum, small molecule antiviral treatments, much of the early COVID-19 therapeutic research focused on the potential repurposing of compounds already approved by the United States Food and Drug Administration (FDA) for other diseases [6–8]. Though an overwhelming majority of compounds did not prove effective, a small number were tested in clinical trials, with only the nucleoside analogues Remdesivir and Molnupiravir receiving FDA Emergency Use Authorization or full approval for the treatment of early COVID-19 by the time of writing [9, 10]. In addition to repurposed compounds, SARS-CoV-2-specific therapies have been authorized or approved by the FDA, including the oral protease inhibitor Paxlovid and several monoclonal antibody treatments [11–14].

SARS-CoV-2 infection begins with binding of the surface protein Spike (S) to the cellular receptor, angiotensin converting enzyme II (ACE2), following which S can be cleaved on the cell surface by Transmembrane protease serine 2 (TMPRSS2) or by Cathepsins B or L in endosomes to facilitate membrane fusion and viral genome release into the cytosol [15–21]. Nonstructural proteins are then immediately translated to a polyprotein to be liberated into individual proteins by viral mediated proteolysis, leading to the formation of replication organelles (DMVs) [22–25]. RNA replication then occurs in these organelles, resulting in the generation of new viral genomes and the production of viral subgenomic RNA (sgRNA), the coronavirus equivalent of mRNA, that are translated into structural and accessory proteins [22, 26]. Assembly of new SARS-CoV-2 virions occurs at the ER-Golgi Intermediate Compartment (ERGIC) followed by virus exit by a still undefined exocytotic method [27–29].

Interestingly, all therapeutics currently authorized or approved by the FDA target early stages of the SARS-CoV-2 life cycle (e.g., entry, RNA replication). However, later stages of the SARS-CoV-2 life cycle are relatively not well understood and, consequently, late-stage inhibitors that could prove more effective or work synergistically with early-stage inhibitors are lacking. We previously identified 2’, 3’-cyclic nucleotide 3’-phosphodiesterase (CNP) as a late-lifecycle-stage SARS-CoV-2 inhibitor in an overexpression screen of 399 interferon stimulated genes (ISGs) [30]. CNP is a lipid-bound protein that is expressed throughout the body, including pulmonary tissues, but expressed at very high levels in myelin [31–33]. Though early biochemical studies identified CNP by its ability to hydrolyze 2’,3’-cyclic nucleotides, as the name suggests, an exact function and a potential cellular substrate have yet to be demonstrated *in vitro* or *in vivo* [34]. However, a growing pool of evidence has begun to establish potential functions for the mitochondria-targeted isoform of CNP (CNP2), which has been shown to inhibit opening of the mitochondrial permeabilization transition pore (mPTP), a process that leads to mitochondrial depolarization in response to mitochondrial Ca^2+^ overload, cellular oxidative stress, or potentially SARS-CoV-2 infection [35–39]. In our previous work, we showed CNP overexpression inhibited SARS-CoV-2 replication in all the cell lines tested, including primary human tracheobronchial epithelial cells, and further showed the inhibition to occur at a replication lifecycle stage post-genome replication [30]. However, the exact lifecycle stage affected and the mechanism of action has yet to be identified.

In this study, we confirm that overexpression of CNP *in vitro* has no effect on SARS-CoV-2 genome replication and further show CNP affects neither subgenomic RNA generation nor structural protein translation. However, CNP overexpression reduced SARS-CoV-2 N localization to sites of virion assembly (ERGIC) and reduced intracellular SARS-CoV-2 titers, providing strong evidence that virion assembly is inhibited. Additionally, we show that CNP inhibition of SARS-CoV-2 required mitochondrial targeting of CNP, which reduced SARS-CoV-2-induced mPTP opening and mitochondrial reactive oxygen species (ROS) release. Indeed, replacing reactive oxygen species in CNP stably-expressing cells through peroxide treatment resulted in a rescue of virus titer. Finally, we show that CNP overexpression in Balb/c laboratory mouse lungs protected mice from SARS-CoV-2 infection, reducing lung virus titers by over 1,000-fold. This work highlights the importance of cytosolic ROS for SARS-CoV-2 virion assembly and identifies CNP as a potential therapeutic target for COVID-19.

## Results

### CNP inhibits SARS-CoV-2 virion assembly

Previous data from our lab shows that CNP overexpression inhibits a SARS-CoV-2 lifecycle stage post-genome replication [30]. To further elucidate the stages affected, HEK293T/hACE2 cells stably expressing HA-tagged CNP or HA-tagged GFP were generated by lentivirus transduction (S1 Fig). Cells were then infected with SARS-CoV-2/WA1 for a 45-minute absorption period followed by washing with cell culture media to remove any remaining unbound virus. Various assays were then used to assess the effects of CNP overexpression on multiple SARS-CoV-2 lifecycle stages. At 6-hours post-infection (hpi), we confirmed that gRNA levels are not significantly different between CNP stable cells and GFP stable cells (Fig 1A) and, further show that sgRNA levels are not altered by CNP (Fig 1B). At 12hpi, western blots for SARS-CoV-2 structural proteins S and N were also not qualitatively different between CNP and GFP stable cells, suggesting no effect on structural protein translation (Fig 1C). However, at 18hpi, SARS-CoV-2 titers were significantly lower when assessed for both intracellular titer (Fig 1D) and extracellular titer (Fig 1E), suggesting the assembly of new SARS-CoV-2 virions is affected by CNP. As SARS-CoV-2 virion assembly takes place at the ERGIC, infected stable cell lines were then fixed at 12hpi and immunofluorescent stained for ERGIC53 and co-stained with either SARS-CoV-2 S or N. Images were captured by confocal microscopy and colocalization was assessed. HEK293T/hACE2 cells stably expressing CNP showed higher colocalization between ERGIC and SARS-CoV-2 S as compared to GFP control cells (Fig 2A, quantified in Fig 2C). However, colocalization between SARS-CoV-2 N and ERGIC markers was markedly lower for CNP expressing cells as compared to GFP controls (Fig 2B, quantified in Fig 2D), suggesting an inhibition of N trafficking to sites of virion assembly.

**Fig 1:**
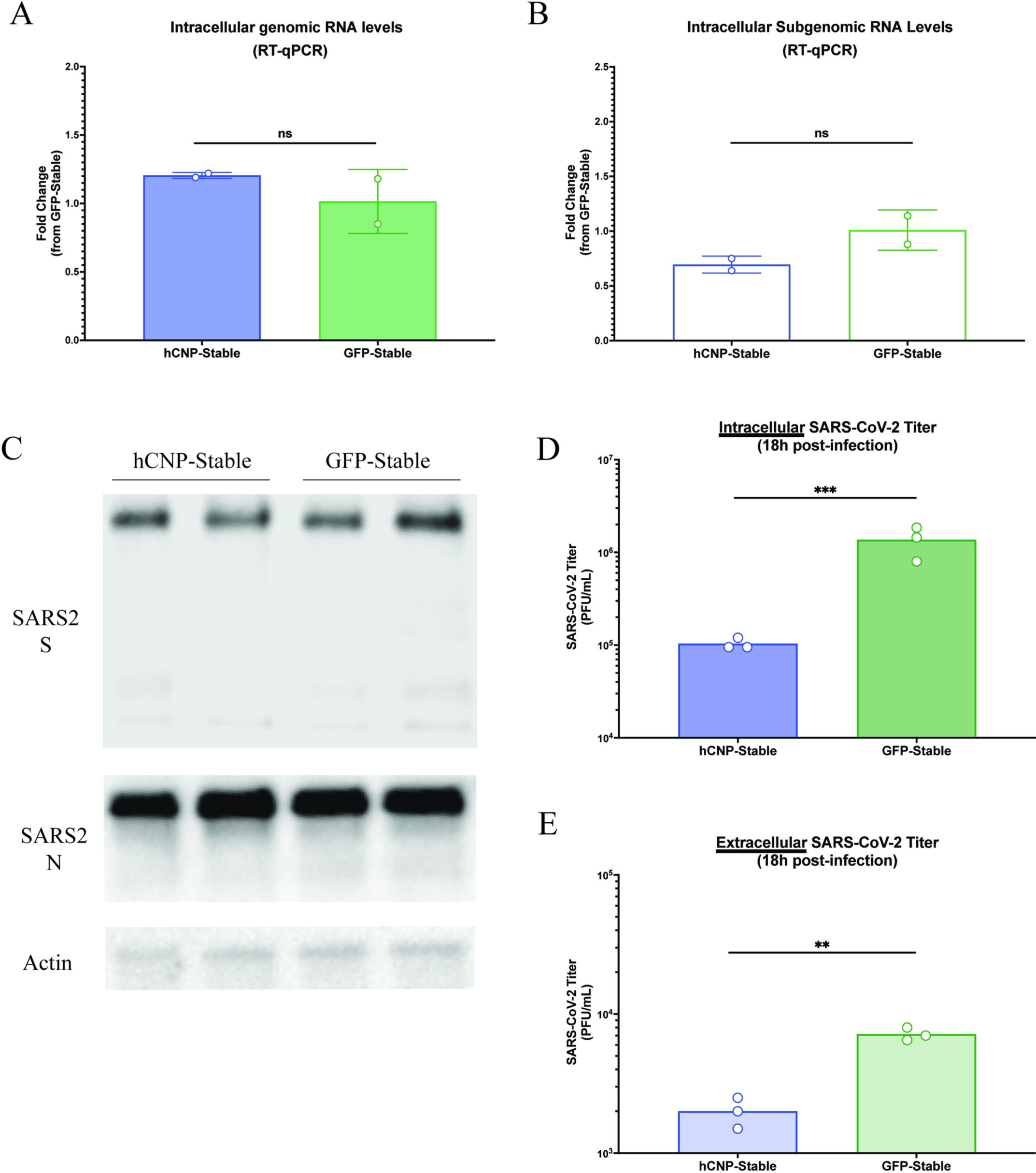
CNP overexpression inhibits generation of new SARS-CoV-2 virions. HEK293T/hACE2 cells stably overexpressing either CNP or GFP were infected with SARS-CoV-2/WA1 for 45-minutes followed by washing of any unbound virus. At 6-hours post-infection, intracellular SARS-CoV-2 RNA levels were determined by RT-qPCR for (**a**) genomic RNA and (**b**) subgenomic RNA. (**c**) At 12-hours post-infection, lysates were collected and western blot staining performed for SARS-CoV-2 S, SARS-CoV-2 N, and actin as a loading control. At 18-hours post-infection, plaque assays were performed to determine (**d**) intracellular SARS-CoV-2 titers and (**e**) extracellular SARS-CoV-2 titers. All experiments were repeated 2-3 times, as noted by the points in the figure. **p ≤ 0.01, ***p ≤ 0.001, “ns”: p > 0.5.

**Fig 2:**
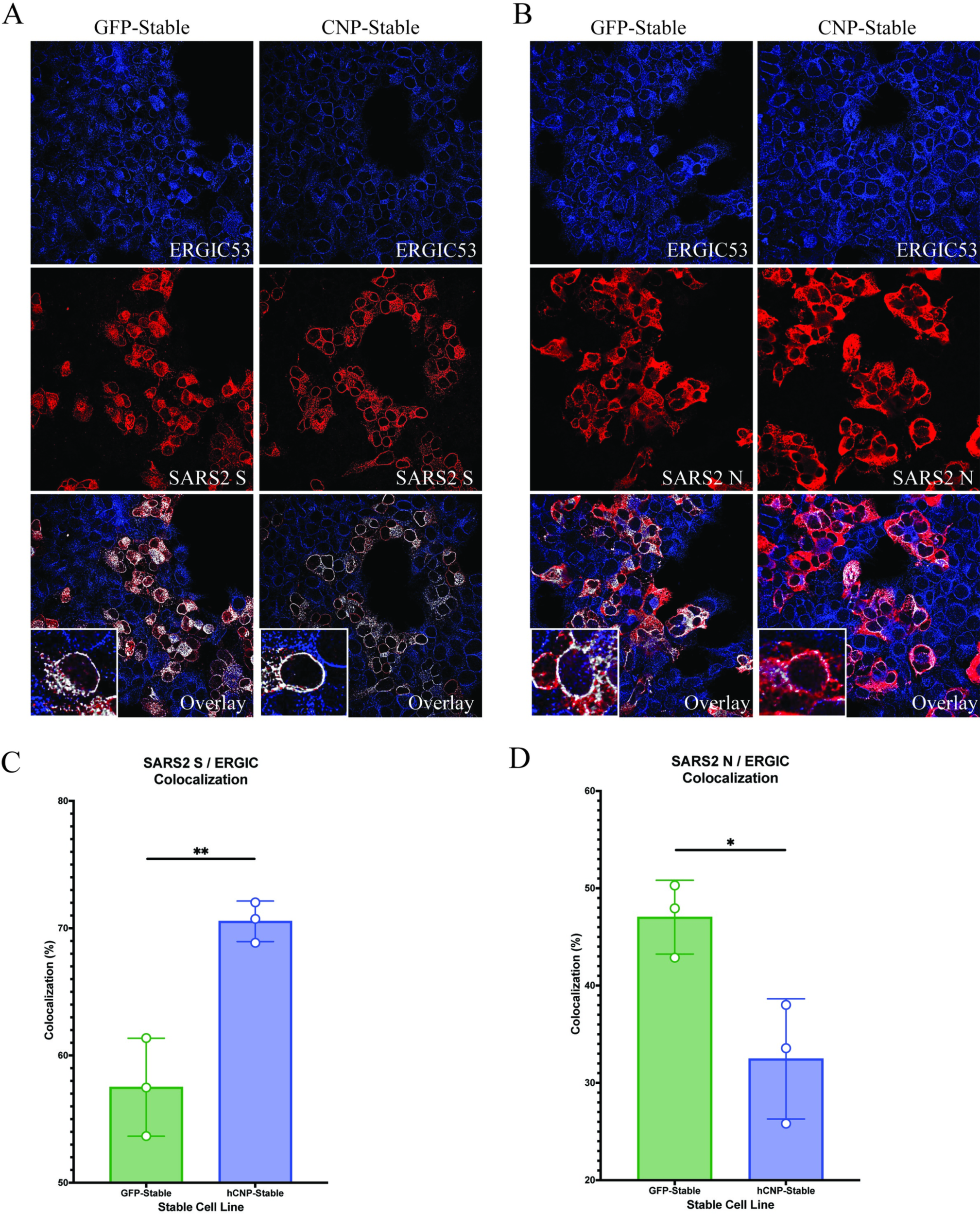
CNP overexpression alters SARS-CoV-2 structural protein localization during virion assembly. HEK293T/hACE2 cells stably overexpressing either CNP or GFP were infected with SARS-CoV-2/WA1 for 45-minutes followed by washing of any unbound virus. At 12-hours post infection, cells were fixed, immunofluorescent stained, and confocal images collected for (**a**) ERGIC53 (blue) and SARS-CoV-2 S (red) or (**b**) ERGIC53 (blue) and SARS-CoV-2 N (red). ERGIC53 colocalization with SARS-CoV-2 structural proteins was assessed in ImageJ for (**c**) SARS-CoV-2 S or (**d**) SARS-CoV-2 N. Higher magnification of each overlay image is shown. *p ≤ 0.1, **p ≤ 0.01. Representative images are shown from three separate replicate experiments.

### CNP inhibits SARS-CoV-2-induced mitochondria depolarization

In order to assess the necessary aspects of CNP required for SARS-CoV-2 inhibition, HA-tagged CNP or HA-tagged CNP mutation/deletion constructs were created, including constructs with mutations to the known catalytic sites (H251A, T253A, H330A, T332A), prenylation site (C418A) and deletions of the mitochondrial targeting sequence (Δ1-20) (S1B Fig, S2 Table). Other domain deletion constructs (i.e., Δ p-loop domain, Δ catalytic domain) were abandoned due to poor expression, likely caused by protein misfolding (S1B Fig). Before analysis of their effect on infection, their localization was assayed in transfected cells. The hCNP wildtype as well as the catalytic site and prenylation site mutant localize to the mitochondria while the MTS deletion mutant is diffuse in the cytoplasm (S1C Fig). To determine the domains of CNP required for SARS-CoV-2 inhibition, HEK293T/hACE2 cells were transfected with HA-tagged wildtype and mutant CNP constructs. The day following transfections, cells were infected with SARS-CoV-2/WA1 (MOI = 0.01) for 48 hours, at which point supernatants were collected and SARS-CoV-2 titers determined by plaque assay. As shown previously, CNP overexpression resulted in a greater than 10-fold decrease in SARS-CoV-2 titers as compared to GFP controls. However, deletion of the mitochondrial targeting sequence resulted in ablation of SARS-CoV-2 inhibition, suggesting targeting of CNP to mitochondria is required for antiviral activity (Fig 3A).

**Fig 3:**
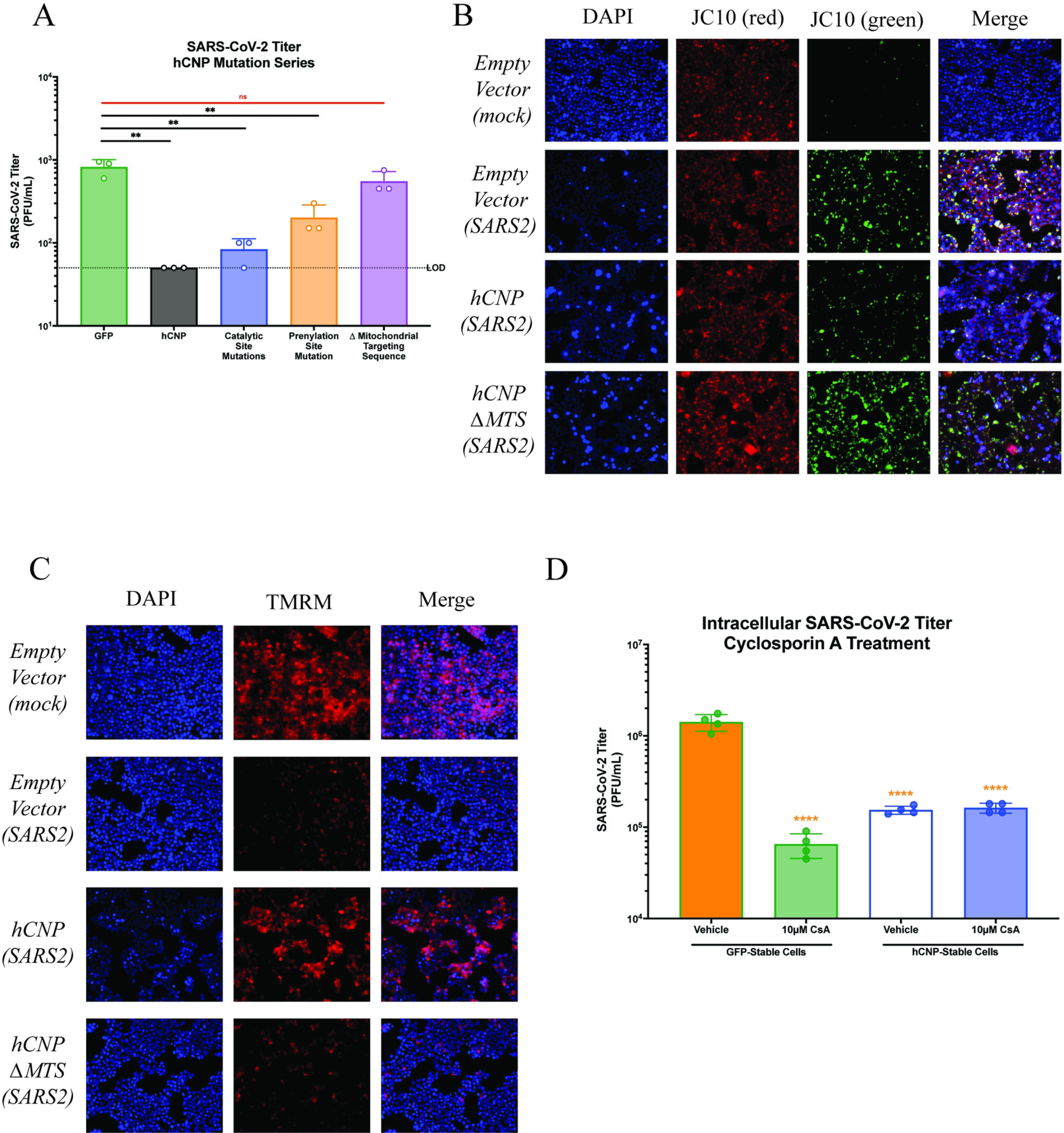
CNP overexpression inhibits SARS-CoV-2-induced mitochondrial depolarization. HEK293T/hACE2 cells were infected with SARS-CoV-2/WA1 24-hours following transfection with indicated plasmids. (**a**) 48-hours post-infection, supernatants were collected, and SARS-CoV-2 titers were determined by plaque assay. (**b**-**c**) At 6-hours post-infection, mitochondrial function was assessed for (**b**) mitochondrial depolarization with JC10 staining or (**c**) opening of the mPTP with TMRM staining. (**d**) HEK293T/hACE2 cells stably overexpressing either CNP or GFP were infected with SARS-CoV-2/WA1 (MOI = 5) for 45-minutes followed by washing of any unbound virus. Fresh media containing 10µM Cyclosporin A or DMSO was then immediately added and intracellular SARS-CoV-2 titer determined at 18hpi. **p ≤ 0.01. Representative images are shown for JC10 and TMRM experiments from three replicates.

Given this result, we aimed to assess the effects of CNP overexpression on mitochondrial function during SARS-CoV-2 infection. Previous work has shown that SARS-CoV-2 infection leads to mPTP-induced mitochondrial depolarization as soon as 3hpi [39]. Additionally, others have shown that CNP inhibits the formation and function of the mPTP [36, 37]. To further assess this relationship, HEK293T/hACE2 cells were transfected with hCNP, hCNPΔMTS, or empty plasmid constructs as described above and infected the following day with SARS-CoV-2/WA1 (MOI = 5). At 6hpi, cells were stained with JC10 dye to assess mitochondrial depolarization or TMRM dye to assess mPTP opening. As was shown previously, SARS-CoV-2 infection in control-transfected cells and hCNPΔMTS-transfected cells led to mitochondrial depolarization, as shown by increases in JC10 staining (Fig 3B) and opening of the mPTP as shown by loss of TMRM staining, as compared to mock infected cells. However, these effects were not observed for hCNP-transfected cells, suggesting hCNP inhibits SARS-CoV-2-mediated mitochondrial depolarization.

This data suggests that CNP could be directly regulating the mPTP opening and release of ROS. While components of the mPTP are not agreed upon, the inhibition of mPTP opening by Cyclosporin A is. We hypothesize that if hCNP is regulating mPTP opening then further inhibition by Cyclosporin A will not cause additional reduction in virus titer. If however CNP is targeting a different process in the mitochondria then the addition of Cyclosporin A before infection will additively or synergistically lead to an increased inhibition of SARS-CoV-2 replication. To further assess the effects of hCNP on mPTP function, HEK293T/hACE2 cells stably expressing either hCNP or GFP were infected with SARS-CoV-2 followed by treatment with 10µM Cyclosporin A, an established inhibitor of mPTP function, or DMSO as a vehicle control. Intracellular SARS-CoV-2 titers were significantly lower when Cyclosporin A treatment was administered to GFP stable cells but not lower for hCNP stable cells, as compared to vehicle controls (Fig 3D). We conclude that this means that Cyclosporin A and hCNP could be targeting the same complex in mitochondria, the mPTP.

### Mitochondrial reactive oxygen species release is needed for efficient SARS-CoV-2 virion assembly

We have confirmed that SARS-CoV-2 infection can induce mitochondrial dysfunction and further show that this dysfunction can be inhibited by hCNP. One major consequence of sustained mitochondrial depolarization is the production and release of ROS. To assess the importance of ROS to SARS-CoV-2 virion assembly inhibition, HEK293T/hACE2 cells were transfected with hCNP, hCNPΔMTS, the catalytic site mutant (hCNP/Cat), the prenylation site mutant (hCNP/Pren) or empty plasmid constructs as described above and infected the following day with SARS-CoV-2/WA1 (MOI = 5). At 6hpi, cells were stained with H_2_DCFDA to visualize cytosolic ROS (Fig 4A). As was shown previously, SARS-CoV-2 infection in control-transfected cells and hCNPΔMTS-transfected cells led to increased ROS production, as compared to mock infected cells. The hCNP/Cat mutant maintains inhibition of ROS production after infection, similar to the wildtype hCNP. The hCNP/Pren mutation has a moderate phenotype. The inhibition of ROS generation reflects the virus inhibition seen in Fig 3. This data suggests that hCNP-inhibition of mitochondrial depolarization may lead to inhibition of ROS production and release into the cytosol.

**Fig 4:**
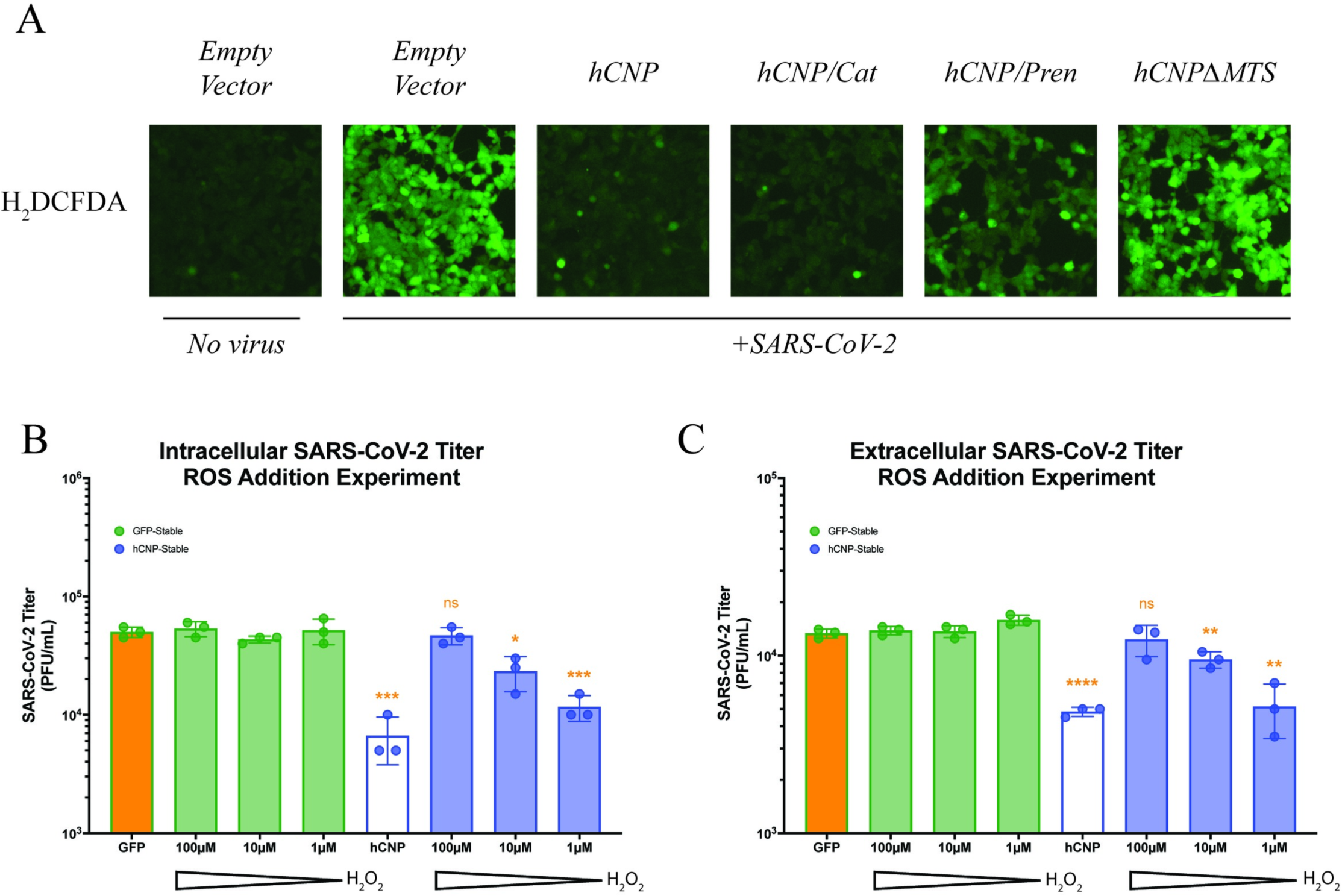
Reactive oxygen species are important for SARS-CoV-2 virion assembly. **(a)** HEK293T/hACE2 cells were infected with SARS-CoV-2/WA1 24-hours following transfection with indicated plasmids. At 6-hours post-infection, the presence of cytosolic reactive oxygen species was assessed by H_2_DCFDA staining. (**b-c**) HEK293T/hACE2 cells stably overexpressing either CNP or GFP were infected with SARS-CoV-2/WA1 (MOI = 5) for 45-minutes followed by washing of any unbound virus. At 3-hours post-infection, cells were treated with a concentration range of hydrogen peroxide. At 18-hours post-infection, plaque assays were performed to determine (**b**) intracellular SARS-CoV-2 titers and (**c**) extracellular SARS-CoV-2 titers. *p ≤ 0.1, **p ≤ 0.01, *** p ≤ 0.001, ****p ≤ 0.0001. Representative images are shown for H_2_DCFDA staining from three replicates. Virus titer data is the shown for three experiments.

To further confirm the importance of ROS production to SARS-CoV-2 replication, we infected HEK293/hACE2 cells stably expressing either hCNP or GFP with SARS-CoV-2/WA1 (MOI = 5) for a 45-minute absorption period followed by washing with cell culture media to remove any remaining unbound virus. At 3hpi, cells were then treated with a sub-lethal concentration range of hydrogen peroxide, one of the ROS released following mitochondrial depolarization, and intracellular/extracellular SARS-CoV-2 titers assessed at 18hpi. The addition of ROS to hCNP-stable HEK293T/hACE2 cells resulted in a dose-dependent rescue of SARS-CoV-2 titers as compared to GFP control cells when assessed for intracellular (Fig 4B) or extracellular (Fig 4C) titers. This data suggests that ROS release from the mitochondria is required for SARS-CoV-2 virion assembly and release.

### CNP reduces SARS-CoV-2 infection in a mouse model of COVID-19

To assess the effects of CNP on SARS-CoV-2 infection in a mouse model of COVID-19, rAd5 vectors were generated to dually express hACE2, the SARS-CoV-2 entry receptor, and either HA-tagged hCNP or HA-tagged eGFP. To validate rAd5 transduction efficiency, Balb/c laboratory mice were transduced intranasally with 1 x 10^8^ viral particles (VP) per mouse and effects on weight change and hCNP and hACE2 expression were assessed 3-days post-transduction (S3A Fig). rAd5 transduction resulted in minimal weight-loss (S3B Fig) by 3-days post-transduction and we observed no qualitative differences in pulmonary inflammation as shown by histology. Transduction by either rAd5 vector resulted in increased expression of hACE2 (Fig 5c) and HA-tagged hCNP or HA-tagged eGFP (S3D Fig) as visualized by immunofluorescent staining of lung histology sections. Additionally, levels of hACE2 mRNA were elevated due to either rAd5 transduction (Fig 5E) and hCNP mRNA was elevated in rAd5/hCNP transduced animals (S3F Fig).

**Fig 5:**
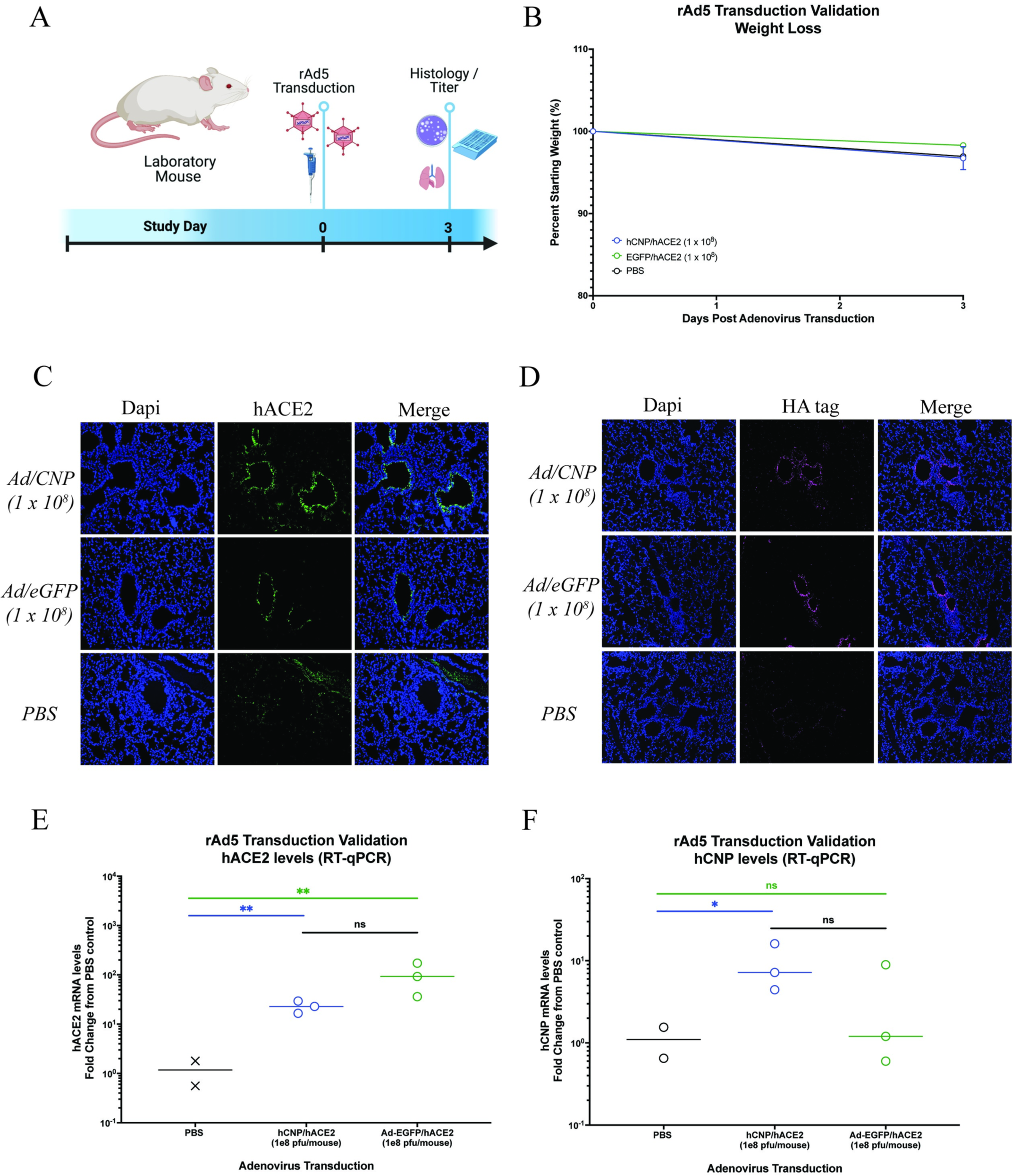
CNP transduction reduces SARS-CoV-2 infection in Balb/c laboratory mice. **(a)** Groups of mice were transduced intranasally with rAd5 vectors (1 x 10^8^ viral particles/mouse) dually expressing hACE2 and either HA-tagged CNP or HA-tagged eGFP 3 days prior to intranasal infection with SARS-CoV-2/WA1 (1 x 10^3^ or 1 x 10^5^ PFU/mouse). Image created using BioRender. **b**-**c** Infectious viral loads from lung homogenates were determined by plaque assay at 2- or 4-days post SARS-CoV-2 challenge for the (**b**) 1 x 10^3^ PFU/mouse or (**c**) 1 x 10^5^ PFU/mouse challenge groups. **d**-**e** Virus titers were also determined by RT-qPCR for sgRNA for the (**d**) 1 x 10^3^ PFU/mouse or (**e**) 1 x 10^5^ PFU/mouse challenge groups. *** p ≤ 0.001, ****p ≤ 0.0001. Virus titer data is the shown for three experiments.

Using these validated rAd5 vectors, we next assessed effects of CNP on SARS-CoV-2 infection in an animal COVID-19 disease model. Balb/c laboratory mice were intranasally transduced with rAd5 vectors (1 x 10^8^ VP/mouse) 3 days-prior to intranasal challenge with SARS-CoV-2/WA1 (1 x 10^3^ pfu/mouse or 1 x 10^5^ pfu/mouse) (Fig 5A). Importantly, the WA1 strain of SARS-CoV-2 cannot enter cells using murine ACE2 and will only be able to infect rAd5-transduced cells expressing hACE2. hCNP overexpression resulted in a greater than 1000-fold decrease in live virus titers to undetectable levels from mouse lung homogenates at 2- or 4-dpi for either the 1 x 10^3^ pfu/mouse (Fig 5B) or 1 x 10^5^ pfu/mouse (Fig 5C) challenge groups. Additionally, there was a marked decrease in SARS-CoV-2 sgRNA in lungs from hCNP transduced animals as compared to control animals for either 1 x 10^3^ pfu/mouse (Fig 5D) or 1 x 10^5^ pfu/mouse (Fig 5E) challenge groups.

## Discussion

Therapeutics against COVID-19 remain limited to early SARS-CoV-2 replication cycle targets, including monoclonal antibody treatments that block entry and nucleoside analogues that inhibit viral RNA replication. This is in part due to the relative gap in understanding that remains for later SARS-CoV-2 lifecycle stages. Our work aimed to further define the mechanism of action of a potential late-stage inhibitor of SARS-CoV-2, CNP, to both assess its therapeutic potential and to further investigate later SARS-CoV-2 lifecycle stages. Consistent with our previously published work, we confirmed that CNP overexpression had no effect on SARS-CoV-2 genome replication but resulted in a significant decrease in extracellular SARS-CoV-2 titers, again suggesting CNP inhibits somewhere between these viral lifecycle steps. We further showed that the generation of sgRNA, the equivalent of the viral mRNA, and translation of SARS-CoV-2 structural proteins were similarly not affected by CNP overexpression. However, intracellular SARS-CoV-2 titers were significantly reduced in CNP overexpressing cells, suggesting the generation of newly made virions is hampered in the presence of CNP. Indeed, CNP overexpression resulted in a reduction of N localization to sites of virion assembly, the ERGIC, and resulted in higher amounts of S localization at the ERGIC, suggesting unincorporated S may be sequestered at the ERGIC awaiting assembly with N. Collectively, these data show that CNP inhibits the assembly of new SARS-CoV-2 virions.

Our work also demonstrates the importance of mitochondrial depolarization to efficient virion assembly. SARS-CoV-2 infection has been shown previously to lead to mitochondrial dysfunction and the release of ROS, though no specific connection to viral replication was made [39]. Our study shows that CNP targeting to mitochondria was essential for SARS-CoV-2 inhibition and that this targeting decreased SARS-CoV-2-induced mitochondrial depolarization and ROS release. Interestingly, the catalytic activity of CNP appeared to be somewhat dispensable for SARS-CoV-2 inhibition, suggesting catalytic activity may not be required for mPTP inhibition, as has been suggested previously [36, 37]. Additionally, the inhibition of ROS production and release caused by mPTP opening appeared to be driving SARS-CoV-2 inhibition, as the addition of ROS to infected CNP-expressing cells by peroxide treatment rescued SARS-CoV-2 intracellular and extracellular titers. We also show that cyclosporin A inhibits SARS-CoV-2 replication to the same level as CNP overexpression and together they show now additional inhibition, suggesting they inhibit the same target. We hypothesize that if they altered different targets then we would see an additive or synergistic effect on viral replication, which we do not. Further research will be needed to assess how ROS release leads to more efficient virion assembly. It is possible that cytosolic ROS may provide a more advantageous environment for post-translational modification to SARS-CoV-2 structural proteins that regulate virion assembly, such as modifications to N that are proposed to regulate N phase separation to promote virion assembly [40]. Alternatively, increases in ROS signaling could promote virion assembly.

This study also shows the potential of CNP to be developed into a potent COVID-19 therapeutic. Our mouse data shows that pre-treatment with CNP inhibited SARS-CoV-2 infection to below detectable levels in lungs throughout the course of infection. Additionally, overexpression of CNP when administered intranasally in mice showed no substantial side effects as measured by weight loss or visualized by histology for pulmonary inflammation, suggesting a CNP treatment could be well tolerated. Additionally, increases in systemic ROS during acute COVID-19 has been observed in humans and implicated in the development of some “Long COVID” symptoms [41]. It is possible that CNP treatment could alleviate some of these long-term effects. However, further development is needed to transition a potential CNP treatment from rAd5 transduction to a more viable treatment option. Alternatively, other mPTP inhibitors, such as Cyclosporin A, could show additional promise as a COVID-19 treatment *in vivo* than what has already been demonstrated *in vitro* [39]. Collectively, this work shows the potential of mPTP inhibition and reduction in ROS production as a treatment for COVID-19 and the potential of CNP to be a new SARS-CoV-2 antiviral target.

## Material and Methods

### Data Availability

All data for this manuscript is available at the Center for Open Science at https://osf.io/f6db3/

### Ethics Statement

All experiments were approved by the University of Maryland at Baltimore, Institutional Biosafety Committee and the Institutional Animal Care & Use Committee. All experiments with live virus were performed in a biosafety level 3 facility at The University of Maryland at Baltimore.

### Cells

VeroE6 cells overexpressing TMPRSS2 were generously provided by Dr. Makoto Takeda (VeroT, [42]). VeroT and HEK293T cells overexpressing hACE2 (HEK293T/hACE2) (ATCC BEI NR-52511) were cultured in DMEM medium (Quality Biological) supplemented with 10% (vol/vol) heat-inactivated FBS (Sigma), 1% (vol/vol) penicillin–streptomycin (Gemini Bio-Products), 1% (vol/vol) l-glutamine (2 mM final concentration; Gibco), and 1% (vol/vol) HEPES buffer (10mM final concentration; Gibco).

### Virus

All work with SARS-CoV-2 was performed in an A/BSL3 laboratory and approved by our Institutional Biosafety Committee (IBC# IBC-00005484) and Institutional Animal Care and Use Committee (IACUC# 1120004). The original strain of SARS-CoV-2 was provided by the CDC following isolation from a patient in Washington State (WA-1; BEI number NR-52281). Formal consent was obtained for sampling via the CDC. Stocks were prepared by infection of VeroT cells for two days when a cytopathic effect was starting to become visible. The media were collected and clarified by centrifugation before being aliquoted for storage at −80 °C. The titer of the stock was determined by plaque assay using VeroT cells.

Lentiviruses and adenoviruses were purchased from and prepared by VectorBuilder Inc. Any infections with lentiviruses and adenoviruses were performed in an A/BSL2 or A/BSL3 laboratory and approved by our Institutional Biosafety Committee (IBC# IBC-00005484) and Institutional Animal Care and Use Committee (IACUC# 1120004).

### Semi-solid Avicel Plaque Assay

Plaque assays were performed similar to what has been described previously [43]. Briefly, 12-well plates were seeded with 2.5 x 10^5^ VeroT cells/well one day prior to processing. On the day of processing, media was removed from the 12-well plates and 200µL of dilutions of sample in DMEM were added to each well. Plates were incubated at 37 °C (5% CO_2_) for 1 hour with rocking every 15 minutes. Following incubation, 2mL of plaque assay media, DMEM containing 0.1% agarose (UltraPure) and 2% FBS (Gibco), was added to each well and incubated for 2 days at 37 °C (5% CO_2_). Following incubation, plates were fixed with 4% paraformaldehyde, stained with 0.25% crystal violet (w/v), plaques counted, and titers calculated as plaque forming units (PFU).

### RT-qPCR

RT-qPCR was processed for cell lysate and for mouse lung homogenates as described previously [43]. RNA was extracted per the manufacturer’s instructions using the Direct-zol RNA Miniprep Kit (Zymo Research). RNA was converted into cDNA (Thermo RevertAid Reverse Transcriptase) and used as template for qPCR (Qiagen RT2 SYBR green qPCR Mastermix). Primers are listed in S1 Table. Data was collected on a 7500 Fast Dx Real-Time PCR Instrument (Applied Biosystems).

### Plasmids

Full length, domain deletion, and functional mutation human CNP gene versions were cloned into a pCAGGS-HA as described previously [44]. Briefly, genes were cloned into pCAGGS-HA vectors on EcoR1/Xma1 sites, resulting in a C-terminal HA epitope tag. Genes or gene fragments were PCR amplified with primers (S2 Table) to allow for In-Fusion gene cloning (Takara Bio) according to the manufacturer’s instructions. For functional mutant clones, primer directed mutagenesis was performed and gene fragments were combined during the In-fusion cloning process.

### SARS-CoV-2 inhibition by CNP deletion/mutation constructs

All CNP-mediated SARS-CoV-2 inhibition analysis was performed using HEK293T/hACE2 cells. HEK293T/hACE2 cells were seeded in 24-well plates pre-coated with 50µg/mL Rat Tail Collagen I (Gibco), according to the manufacturer’s instructions. At the time of seeding, pCAGGS-HA plasmids were transfected using 1µL TransIT 2020 (Mirus) per 100µg plasmid and incubated overnight. Cells were then infected with SARS-CoV-2/WA1 (MOI = 0.01). Supernatants were collected 48-hours post-infection and SARS-CoV-2 titers determined by VeroT plaque assay.

### Stable Cell Line Generation

Lentiviruses expressing HA-tagged CNP or HA-tagged GFP were obtained from VectorBuilder, Inc. HEK293T/hACE2 cells were transduced with lentiviruses (MOI = 3) pre-treated with 10µg/mL Polybrene (VectorBuilder, Inc.). Transduced cells were selected by resistance to 10µg/mL Blasticidin (InvivoGen) for 10 days, followed by continued maintenance in cell culture media supplemented with 10µg/mL Blasticidin (InvivoGen).

### SARS-CoV-2 Lifecycle Inhibition Analysis

HEK293T/hACE2 cells stably expressing CNP/HA or GFP/HA were seeded in 24-well plates pre-coated with 50µg/mL Rat Tail Collagen I (Gibco), according to the manufacturer’s instructions, or in Tab-Tek II CC2 chamber slides (Thermo Fisher Scientific). Separate wells were seeded for each assay to process. The following day, cells were infected with SARS-CoV-2/WA1 (MOI = 5) for a 45-minute absorption period, followed by inoculum removal and washing with fresh cell culture media. Cells were then maintained at cell culture conditions until collection.

At 6-hours post infection, cells were washed with complete cell culture media and then lysed in TRIzol (Ambion). Lysates were stored at -80°C until processing for RT-qPCR.

At 12-hours post-infection, cells were washed with complete cell culture media and then lysed in complete lysis buffer containing 20mM Tris-HCl, 150mM NaCl, 1% Nonidet P-40 (v/v), 0.5% Sodium Dodecyl Sulphate (w/v), 5mM EDTA, and 1 X complete protease inhibitor (Roche), for 5 minutes at room temperature. Lysates were collected and then boiled twice at 95°C for 30 minutes. Lysates were then centrifuged at 21,000 x g for 20 minutes followed by supernatant collection. Supernatants were and stored at -80°C until ready to process by Western Blot (below).

At 12-hours post-infection, cells were fixed in 2% paraformaldehyde for 1 hour at 4°C and then washes once with 1 x PBS. Fixed cells were stored at 4°C until processing for immunofluorescent microscopy and confocal microscopy.

At 18-hours post-infection, supernatants were collected for extracellular SARS-CoV-2 titer determination and stored at -80°C until VeroT plaque assay processing. Cells were then washed once with complete cell culture media followed by addition of 120µL of complete cell culture media. Cells were then freeze-thawed three times to burst cells and lysates were collected for intracellular SARS-CoV-2 titer determination and stored at -80°C until VeroT plaque assay processing.

### Western Blot

Western blots were performed similar to what has been described previously [45]. Cell lysates were run on Mini-PROTEAN TGX Precast Gels (BioRad) and tank blotted onto PVDF membranes according to the manufacturer’s instructions. Following blotting, membranes were blocked in blocking buffer, TBST (TBS (from where?) with 0.1% Tween 20 (v/v) (from where)) and 5% milk (w/v) (from where), for 1 hour at room temperature. Membranes were then incubated in primary antibody overnight at 4°C followed by 3 x 10-minute washes with TBST. Membranes were then incubated with diluted secondary antibodies for 1 hour at room temperature followed by 5 x 10-minute washes with TBST. Membranes were then developed with ECL (Cytiva) and imaged with a CCD camera. All antibodies (S3 Table) were diluted in blocking buffer.

### Immunofluorescence Microscopy and Confocal Microscopy

Paraformaldehyde fixed cells were permeabilized in PBS with 0.3% Triton-100 for 30 minutes at room temperature. Cells were then blocked in blocking buffer, PBS containing 5% BSA (w/v), for 1 hour at room temperature. Cells were then incubated with diluted primary antibody overnight at 4°C followed by 3 x 10-minute washes with wash buffer, PBS containing 1% BSA (w/v) and 0.1% Tween-20 (v/v). Cells were then incubated with diluted secondary antibodies for 1 hour at room temperature followed by 3 x 10-minute washes with wash buffer and 2 x 10-minute washes in PBS. Coverslips were then mounted with ProLong Glass Antifade Mountant with NucBlue (Invitrogen) for at least 30 minutes at room temperature. All antibodies (S3 Table) were diluted in blocking buffer.

Stained slides were visualized using a Revolve fluorescent microscope (Echo) or a W-1 Spinning Disk confocal microscope (Nikon). Images collected by confocal microscopy were 2-D deconvoluted using the W-1 spinning disk software and analyzed with ImageJ (NIH) using the Colocalization Threshold colocalization plugin. At least 100 cells per condition were analyzed for colocalization determination in three separate image fields and the percent intensity above threshold colocalized for SARS-CoV-2 S or N with ERGIC53 are reported. Data are representative of two independent experiments.

### Mitochondrial Function Assay Design

HEK293T/hACE2 cells were seeded in Tab-Tek II CC2 chamber slides (Thermo Fisher Scientific). At the time of seeding, pCAGGS-HA plasmids were transfected using 1µL TransIT 2020 (Mirus) per 100µg plasmid and maintained at cell culture conditions overnight. Cells were then infected with SARS-CoV-2/WA1 (MOI = 0.1). Mitochondrial function assays (described below) were then processed at 6-hours post-infection.

### JC-10 Staining

At 6-hours post-infection, JC-10 staining was completed using the mitochondria membrane potential kit (Sigma-Aldrich) according to the manufacturer’s instructions. Slides were then washed once with 1 x HBBS and fixed in 2% paraformaldehyde for 30 minutes at room temperature. Coverslips were then mounted with ProLong Glass Antifade Mountant with NucBlue (Invitrogen) for 30 minutes at room temperature and visualized immediately using a Revolve fluorescent microscope (Echo).

### TMRM Staining

At 6-hours post-infection, opening of the mitochondrial permeabilization transition pore (mPTP) was visualized by tetramethylrhodamine methyl ester perchlorate (TMRM) staining (Sigma). Cells were washed once with 1 x HBBS (Gibco) followed by incubation with 100 nM TMRM diluted in 1 x HBBS for 30 minutes at 37°C. Slides were then washed once with 1 x PBS and fixed in 2% paraformaldehyde for 30 minutes at room temperature. Coverslips were then mounted with ProLong Glass Antifade Mountant with NucBlue (Invitrogen) for 30 minutes at room temperature and visualized immediately using a Revolve fluorescent microscope (Echo).

### Reactive Oxygen Species Staining (H2DCFDA)

At 6-hours post-infection, intracellular reactive oxygen species levels were visualized by H_2_DCFDA staining (Invitrogen) according to the manufacturer’s instructions. Briefly, cells were washed twice with 1 x HBBS (Gibco) followed by incubation with 2.5µM H_2_DCFDA diluted in 1 x HBBS for 30 minutes at 37°C. Slides were then washed three times with 1 x PBS and fixed in 2% paraformaldehyde for 30 minutes at room temperature. Cells were visualized immediately using a Revolve fluorescent microscope (Echo).

### Reactive oxygen species Addition and Cyclosporin-A Treatment Experiments

HEK293T/hACE2 cells stably expressing either CNP/HA or GFP/HA were seeded and infected with SARS-CoV-2/WA1 as described in “SARS-CoV-2 Lifecycle Inhibition Analysis”. At 3-hours post-infection, a concentration range of hydrogen peroxide (Sigma Aldrich) or vehicle (water) was added to infected cells. At 18-hours post infection, supernatants and cell lysates were collected for extracellular/intracellular SARS-CoV-2 titer determination as described in “SARS-CoV-2 Lifecycle Inhibition Analysis” and stored at -80°C until further processing.

For Cyclosporin A treatment studies, HEK293T/hACE2 cells stably expressing either CNP/HA or GFP/HA were seeded and infected with SARS-CoV-2/WA1 as described in “SARS-CoV-2 Lifecycle Inhibition Analysis”. After washing away unbound virus, fresh media containing 10µM Cyclosporin A (SelleckChem) or 0.01% DMSO as a vehicle control was added to cells. At 18-hours post infection, cell lysates were collected for intracellular SARS-CoV-2 titer determination as described in “SARS-CoV-2 Lifecycle Inhibition Analysis” and stored at -80°C until further processing.

### CNP overexpression antiviral Testing *in vivo*

Two mammalian gene expression recombinant Adenovirus serotype 5 (rAd5) viral vectors were prepared by and obtained from VectorBuilder Inc. to dually express hACE2 and either: (1) HA-tagged hCNP (Ad/CNP); or (2) HA-tagged enhanced green fluorescent protein (Ad/eGFP). To confirm rAd5 transduction, 8-10 week old BALB/c laboratory mice were anaesthetized with a mix of xylazine and ketamine diluted in phosphate buffered saline prior to intranasal transduction with 1 x 10^8^ PFU of either Ad/CNP or Ad/eGFP and weights were recorded (Fig 5A). At 3-days post-rAd5-transduction, weights were again recorded and lungs harvested and sectioned for the following: (1) fixed in 4% paraformaldehyde for histopathological analysis and immunofluorescent staining; and (2) homogenized in TRIzol for RT-qPCR analysis. Homogenization occurred using 1.0 mm glass beads (Sigma Aldrich) and a Beadruptor (Omni International Inc.).

For SARS-CoV-2 antiviral testing, 8-10 week old BALB/c laboratory mice were anaesthetized with a mix of xylazine and ketamine diluted in phosphate buffered saline prior to intranasal transduction with 1 x 10^8^ PFU of either Ad/CNP or Ad/eGFP. Three days later, mice were again anaesthetized with a mix of xylazine and ketamine diluted in phosphate buffered saline prior to intranasal inoculation with either 1 x 10^3^ PFU or 1 x 10^5^ PFU of SARS-CoV-2/WA1 (Fig 6A). Control mice were also mock infected with 1 x PBS. Weights were collected daily following SARS-CoV-2 infection and 5 mice sacrificed on 2 dpi and 4 dpi for each treatment group. Lungs were harvested and sectioned for the following: (1) fixed in 4% paraformaldehyde for histopathological analysis; (2) homogenized in TRIzol for RT-qPCR analysis; and (3) homogenized in PBS for plaque assay processing. Homogenization occurred using 1.0 mm glass beads (Sigma Aldrich) and a Beadruptor (Omni International Inc.).

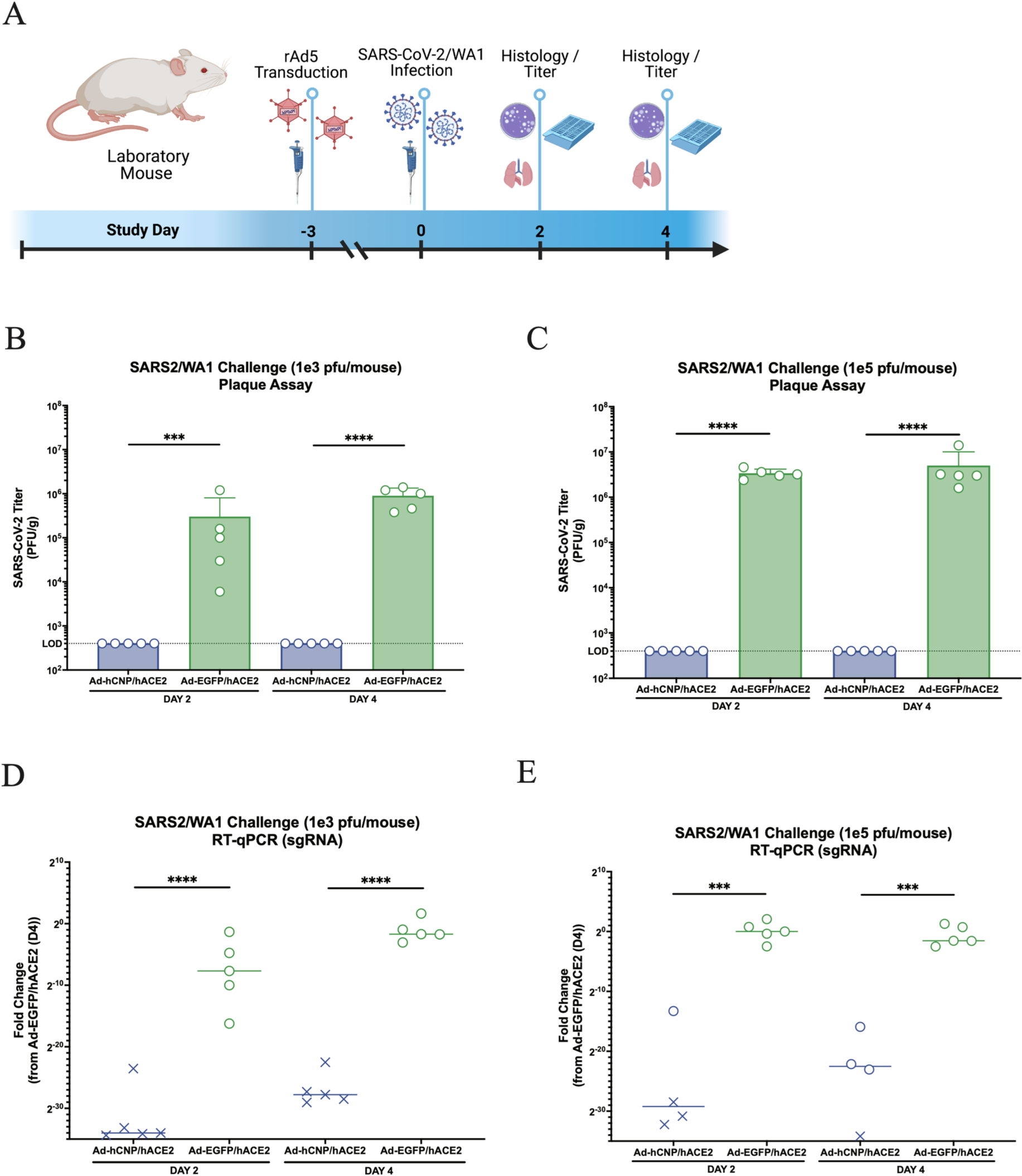

### Histopathology

Histopathology was processed as described previously [46]. Lung sections were fixed in 4% paraformaldehyde in phosphate-buffered saline for a minimum of 48 h, after which they were sent to the Histology Core at the University of Maryland, Baltimore, for paraffin embedding and sectioning. Five-micrometer sections were prepared and used for hematoxylin and eosin (H&E) staining by the Histology Core Services. Sections were imaged at 10x magnification and assembled into figures using Adobe Photoshop and Illustrator software.

### Immunofluorescent staining of mouse lung histology slides

Lung sections were fixed in 4% paraformaldehyde in phosphate-buffered saline for a minimum of 48 h, after which they were sent to the Histology Core at the University of Maryland, Baltimore, for paraffin embedding and sectioning. Slides were then deparaffinized with 2 x 5-minute washes in Xylenes (Sigma Aldrich) followed by rehydration with decreasing concentrations of ethanol in water. Following rehydration, antigen retrieval was completed by boiling samples in citrate buffer (Millipore Sigma) for 5 minutes followed with removal of heat and an additional 15-minute incubation. Samples were then washed twice with ddH2O.

Slides were then stained for immunofluorescent imaging. All incubations were performed in a humidified chamber. Slides were blocked in blocking buffer, PBS++ (1 x PBS with 1mM MgCl and 1mM CaCl_2_) with 1% (w/v) BSA, for 1 hour at room temperature. Following blocking, slides were incubated overnight at 4°C with diluted primary antibody followed by 3 x 10-minute washes with PBS++. Slides were then incubated for 1 hour at room temperature with diluted secondary antibodies, followed by 3 x 10-minute washes with PBS++. Coverslips were then mounted with ProLong Glass Antifade Mountant with NucBlue (Invitrogen) for at least 30 minutes at room temperature and visualized using a Revolve fluorescent microscope (Echo). All antibodies (S3 Table) were diluted in blocking buffer.

## Acknowledgements

The authors would like to thank the Histology Core at the University of Maryland, Baltimore for histology processing.

## Funding

This work is partially funded by NIH R01 AI148166 (to MBF). The funders had no role in study design, data collection and analysis, decision to publish, or preparation of the manuscript.

## Contributions

Conceptualization: J.L., M.B.F. Investigation: J.L., V.M.M., J.A., and M.B.F. Writing – Original Draft Preparation: J.L., V.M.M, and J.A. Writing – Review & Editing: J.L., V.M.M., J.A., and M.B.F.

## Competing Interests Statement

M.B.F. is on the scientific advisory board of Aikido Pharma and has collaborative research agreements with Novavax, AstraZeneca, Regeneron, and Irazu Bio. These do not have any effect on the planning or interpretations of the work presented in this manuscript.

## Supplemental Table / Figures

**S1 Table:**
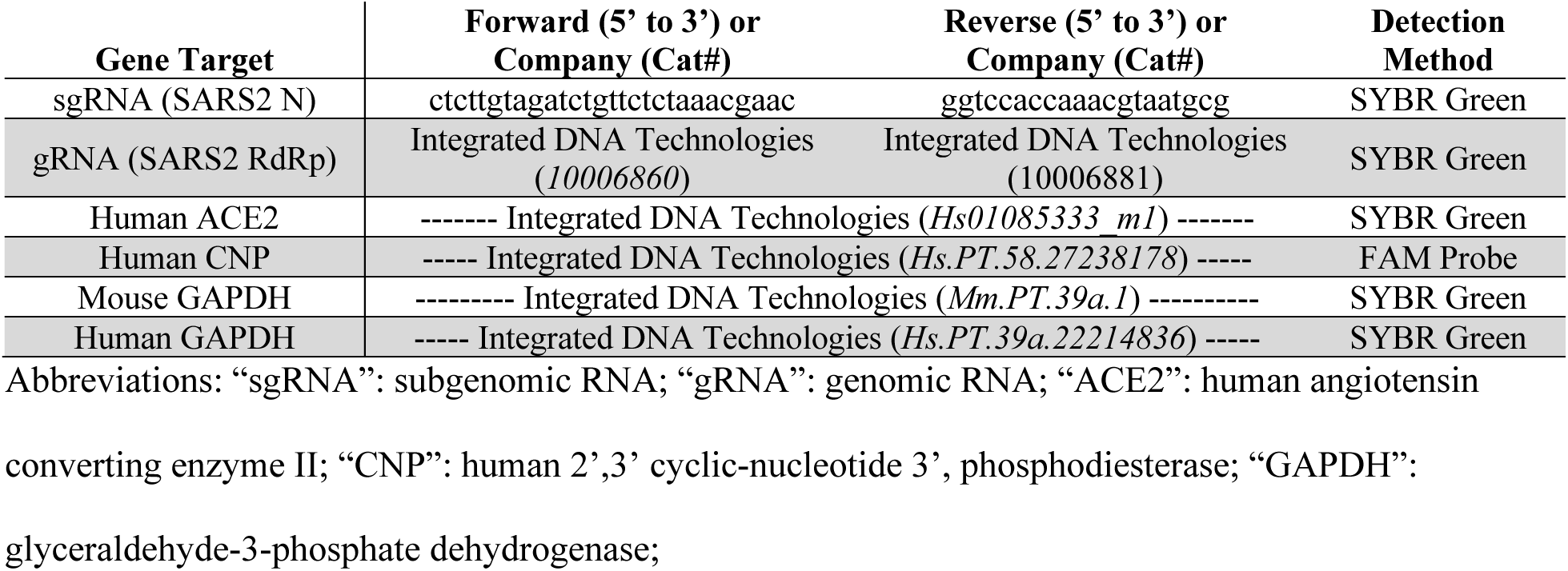
RT-qPCR primers.

**S2 Table:**
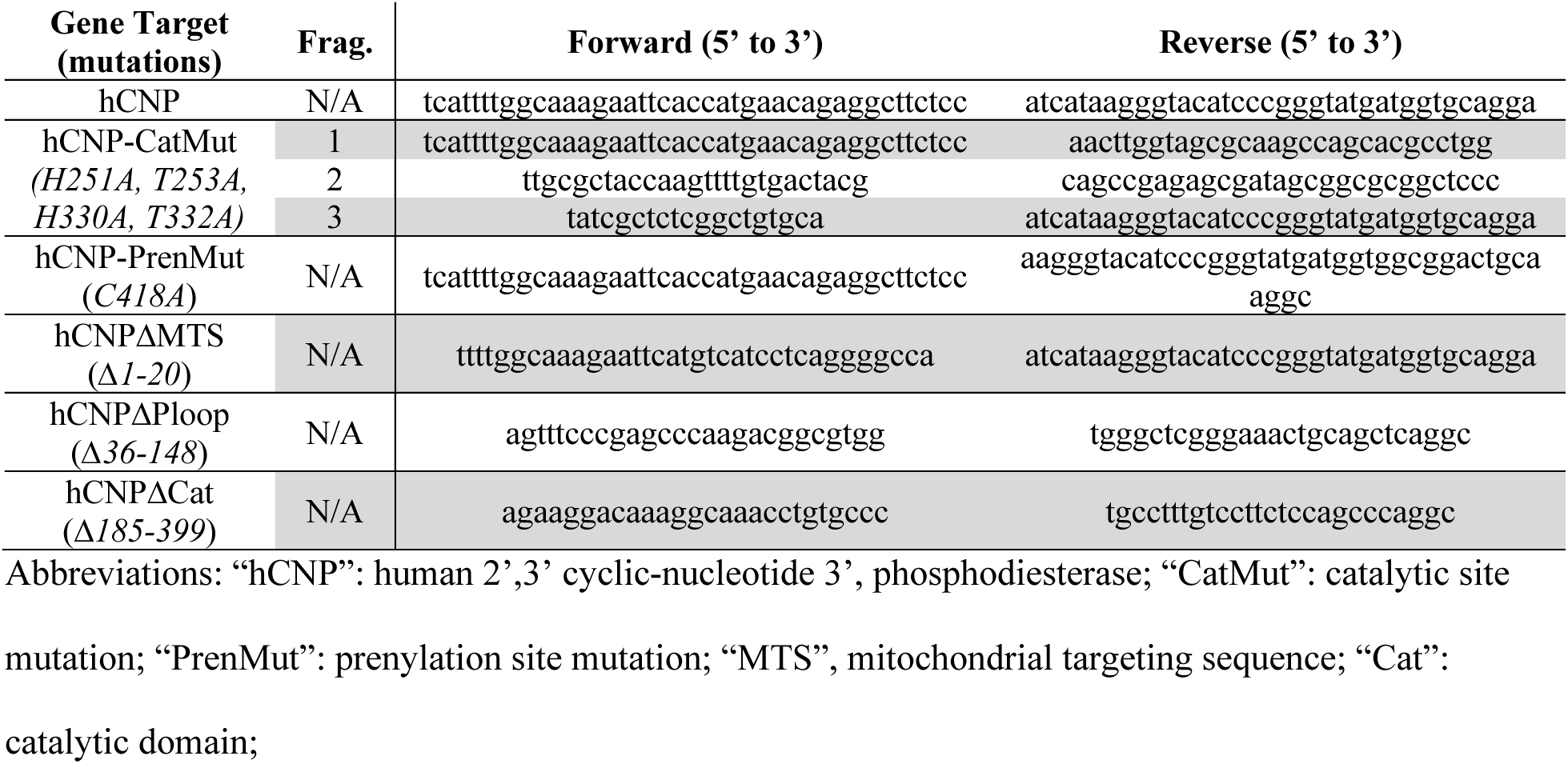
Infusion cloning primers.

**S3 Table:**
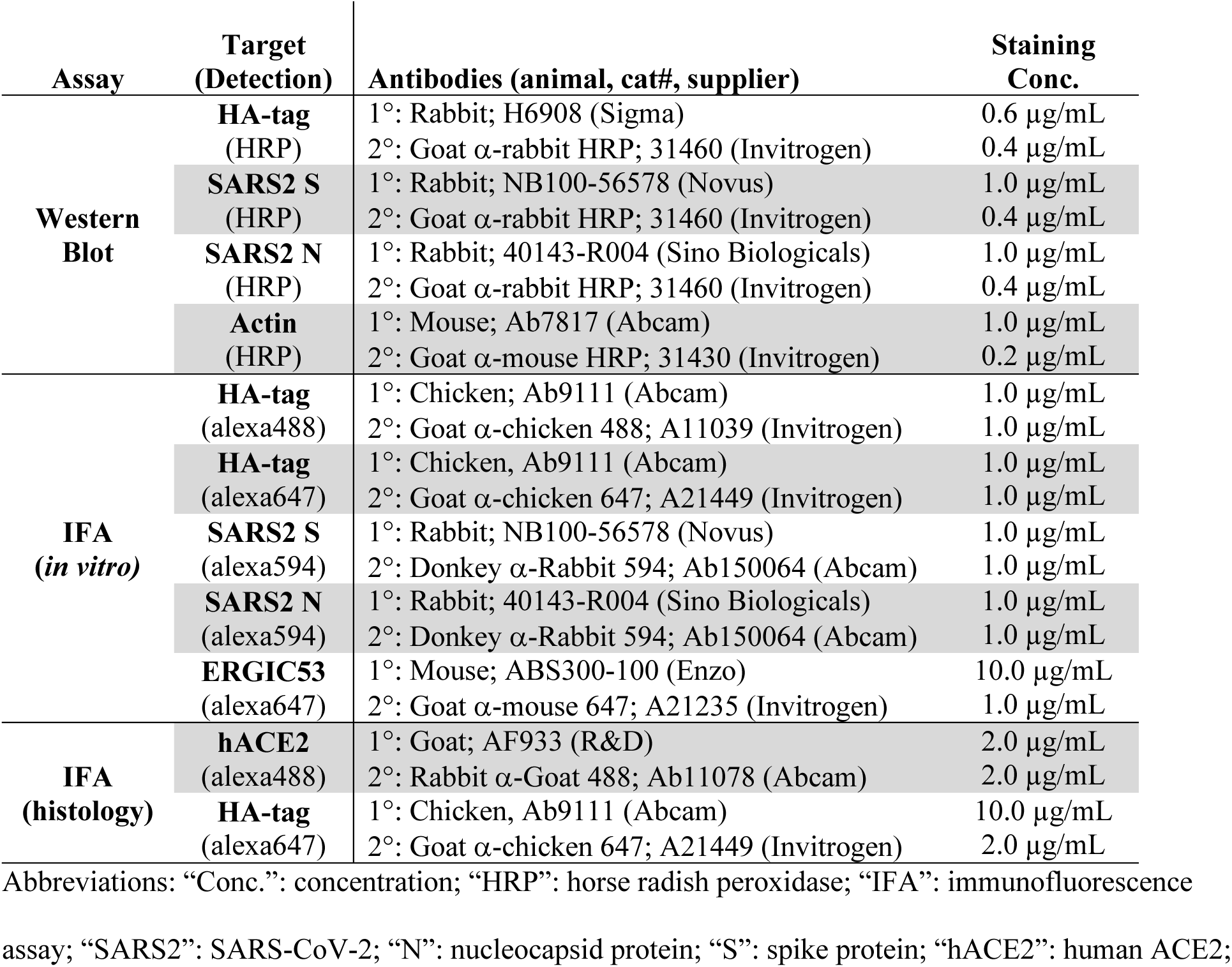
Antibodies for *in vitro* Western Blot and Immunofluorescent staining.

**S1 Fig:**
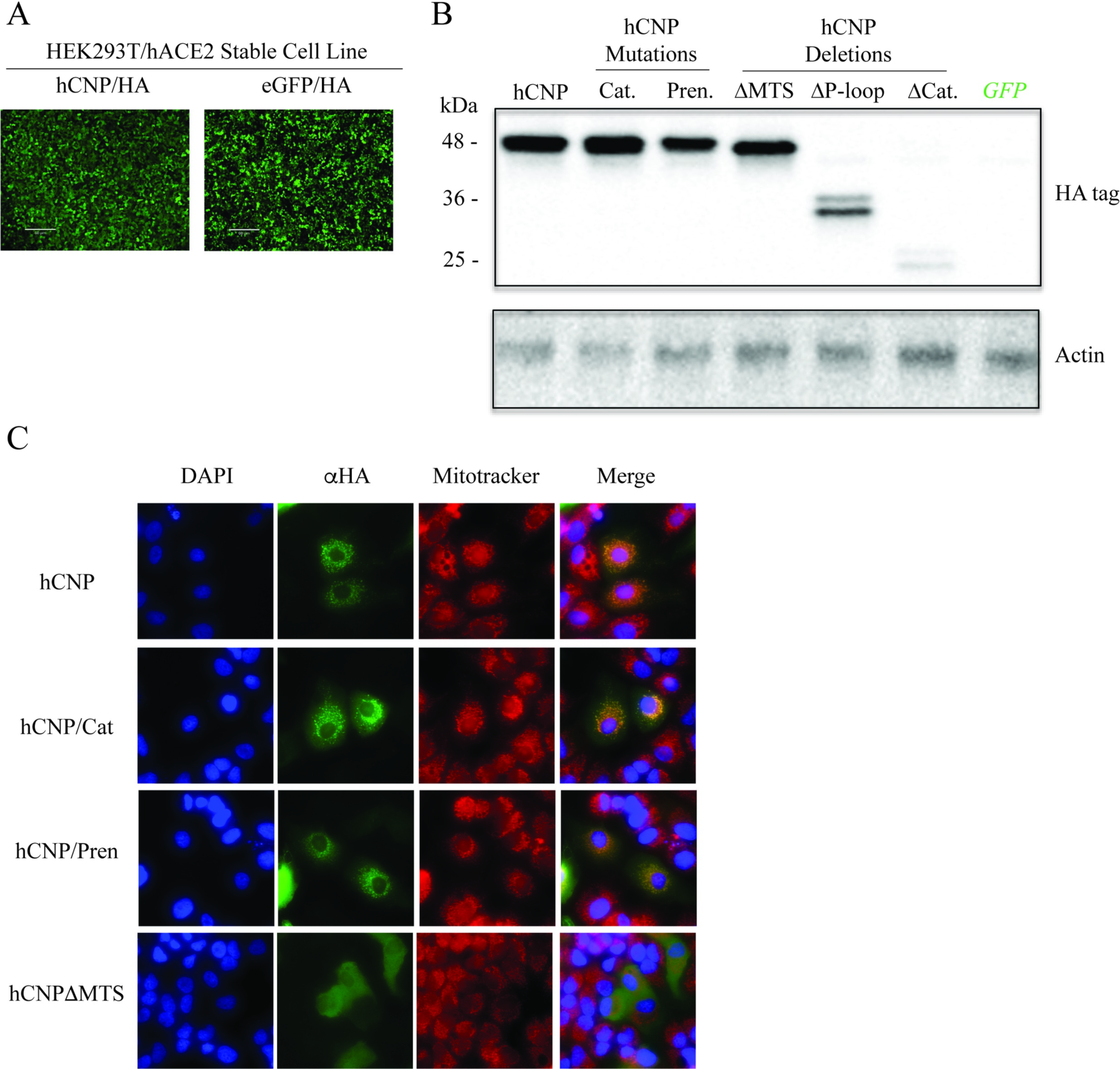
CNP overexpression validation. **(a)** HA-tag staining of HEK293T/hACE2 cells transduced with lentivirus vectors expressing either HA-tagged CNP or HA-tagged eGFP and maintained under selection media. (**b**) Western blots from HEK293T/hACE2 cells transfected with CNP mutation and deletion plasmid constructs stained for HA-tag or Actin controls. (c) Colocalization of HA-tagged CNP constructs in A549/hACE2 cells with mitochondrial staining by mitotracker. Merged images show DAPI, anti-HA and mitotracker colocalization. Image is representative of triplicate samples.

**S2 Fig:**
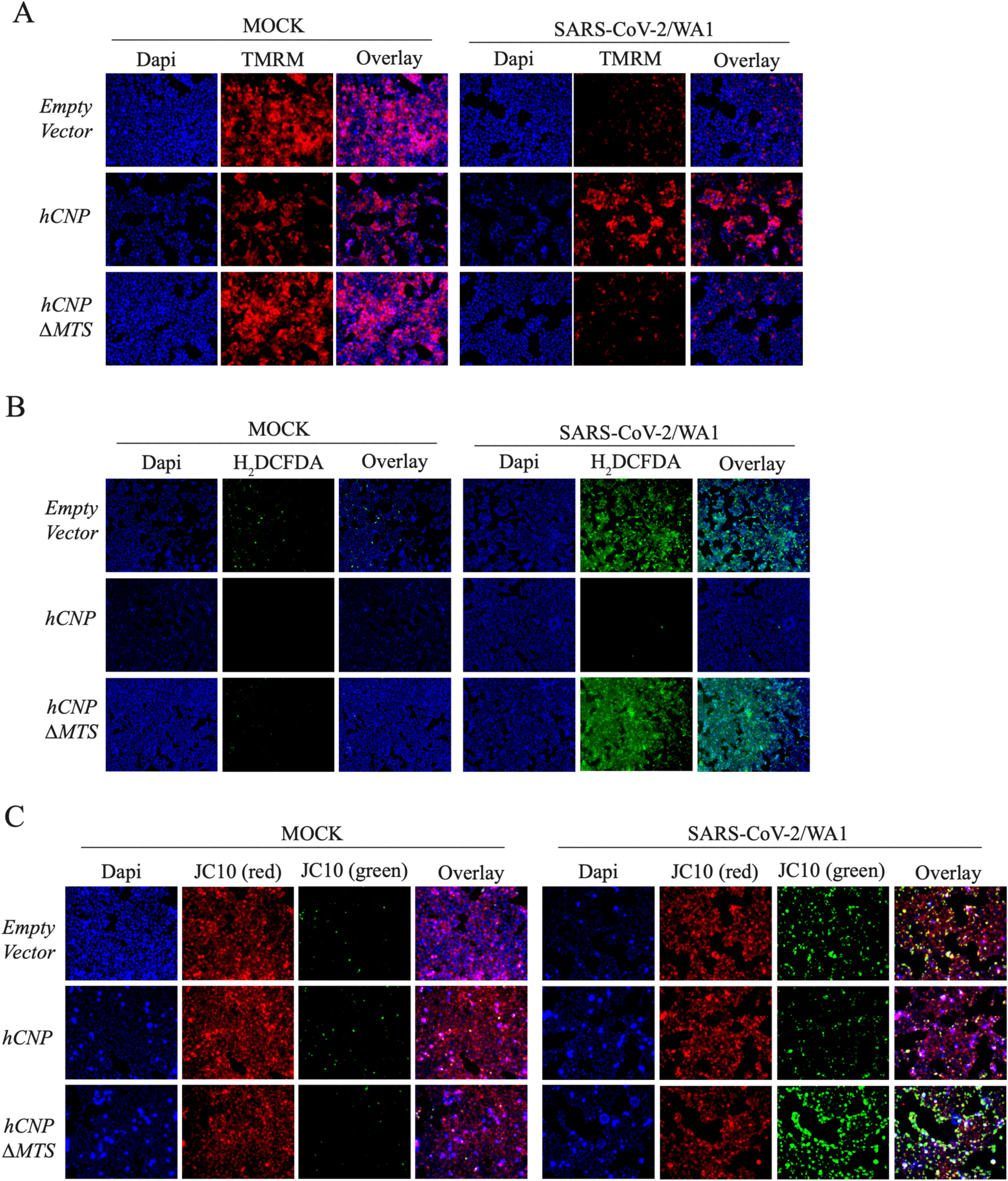
Mitochondrial Function Staining with MOCK infection images. Images correspond to Fig 3. HEK293T/hACE2 cells were infected with SARS-CoV-2/WA1 or MOCK infected 24-hours following transfection with indicated plasmids. At 6-hours post-infection, mitochondrial function was assessed for (**a**) opening of the mPTP with TMRM staining, (**b**) reactive oxygen species staining with H_2_DCFDA, or (**c**) mitochondrial depolarization with JC10 staining.

**S3 Fig:**
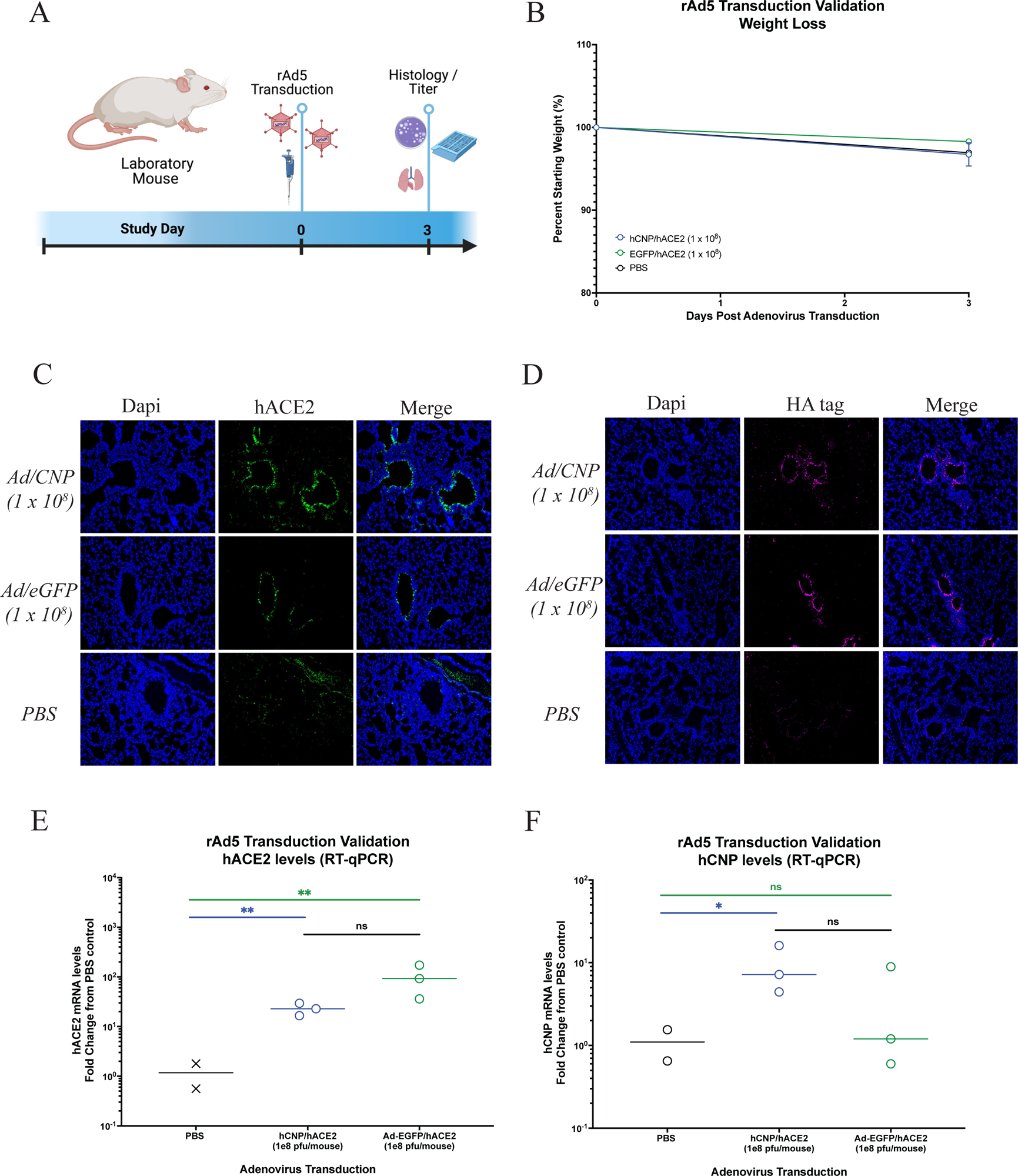
Validation of dual expressing adenovirus vectors in Balb/c laboratory mice. **(a)** Groups of mice were transduced intranasally with rAd5 vectors (1 x 10^8^ viral particles per mouse) dually expressing hACE2 and either HA-tagged CNP or HA-tagged eGFP. Image created using BioRender. (**b**) Weight changes were determined at 3 days post-transduction, plotted as the group mean with error bars indicating the ±SD. **c**-**d** Lungs were collected at 3-days post-transduction and co-stained by immunofluorescence staining for (**c**) nucleus (blue) and hACE2 (green) or (**d**) nucleus (blue) and HA-tag (purple). **e**-**f** RT-qPCR was performed for lung homogenates collected at 3-days post-transduction for (**e**) hACE2 and (**f**) CNP. *p ≤ 0.1, **p ≤ 0.01.

## Notes

### Summary of Updates

Figures and manuscript text have been updated for resubmission.

